# Differential expression between African-ancestry and White patients diagnosed with Triple-Negative Breast Cancer: EGFR, Myc, Bcl2 and β-Catenin as ancestry-associated markers

**DOI:** 10.1101/2020.11.13.381608

**Authors:** Ana T. Matias, Ana Jacinta-Fernandes, Ana-Teresa Maia, Sofia Braga, António Jacinto, M. Guadalupe Cabral, Patrícia H. Brito

**Affiliations:** iNOVA4Health, CEDOC, NOVA Medical School, NMS, Universidade Nova de Lisboa, Lisbon, Portugal; Centre for Biomedical Research, CBMR, Universidade do Algarve, Faro, Portugal; Algarve Biomedical Center, ABC, Universidade do Algarve, Faro, Portugal; Department of Biomedical Sciences and Medicine, DCBM, Universidade do Algarve, Faro, Portugal; Hospital Professor Doutor Fernando Fonseca, HFF, Amadora, Portugal; Instituto CUF de Oncologia, Lisbon, Portugal; UCIBIO, Departamento de Ciências da Vida, Faculdade de Ciências e Tecnologia, FCT, Universidade Nova de Lisboa, Caparica, Portugal; Nova Medical School, NMS, Universidade Nova de Lisboa, Lisboa, Portugal

**Author notes:** Corresponding Authors: Patrícia H. Brito and M. Guadalupe Cabral.

**Keywords:** Triple-negative breast cancer, Ancestry-associated disease risk, African ancestry, Differential gene expression, RNA-sequencing data

## Abstract

**Purpose:** Triple-negative breast cancer (TNBC) has a higher incidence, a younger age of onset, and a more aggressive behavior in African-ancestry women. Biological disparities have been suggested as an important factor influencing the ancestry-associated TNBC discrepancy. In this study, we sought to identify ancestry-associated differential gene and protein expression between African-ancestry and White TNBC patients, controlling for patients’ menopause status and pathological staging at diagnosis.

**Methods:** Differential gene expression analyses (DGEA) were performed using RNA-sequencing data from The Cancer Genome Atlas (TCGA). Gene set enrichment analysis (GSEA) and Ingenuity Pathway Analysis (IPA), with focus on network design, were performed to highlight candidate genes for further validation through immunohistochemistry of TNBC samples from patients followed in Portugal.

**Results:** With 52 African-American and 90 White TNBC patients included, TCGA’s data corroborate that African-American patients have a higher TNBC incidence (28.42% vs 11.89%, p<0.0001). Particularly, premenopausal and stage II disease African-American patients also have significantly lower survival probability, comparing with White patients (log-rank p=0.019 and 0.0038, respectively). DGEA results suggest that expression profile differences are more associated with TNBC staging than with patient’s menopause status. Hippo pathway and cellular community gene sets are downregulated, while breast cancer gene set is upregulated in African-Americans, comparing with White TNBC patients. Furthermore, MAPK pathway gene set is upregulated when controlling for stage II disease. Due to their central role in highly scored networks resulted from IPA’s network design, EGFR, Myc and Bcl2 genes were selected for further validation through immunohistochemistry. We also included β-Catenin in the validation study as it is consensually reported to be required in TNBC tumorigenesis. Although patients used in the DGEA and in the immunohistochemistry experiments are geographically and culturally distinct, both groups of African-ancestry patients are mostly of western-African ancestry and, interesting, differential gene and protein expression matched.

**Conclusions:** We found ancestry-associated gene expression patterns between African-ancestry and White TNBCs, particularly when controlling for menopause status or staging. EGFR, Myc, Bcl2 and β-catenin gene and protein differential expression matching results in distinct populations suggest these markers as being important indicators of TNBC’s ancestry-associated development.

## Introduction

Triple-negative (TNBC) is the breast cancer (BC) subtype that neither expresses estrogen receptor, nor progesterone receptor, nor has Human Epidermal growth factor Receptor 2 (HER2) amplification. With no approved targeted therapies thus far and an aggressive behavior, presenting large locally advanced breast lesion or metastatic disease developing shortly after adjuvant chemotherapy [1, 2], TNBC patients’ survival rate is dismal [3, 4]. Globally, TNBC accounts for 10-20% of invasive BC diagnoses [5–8]. However, TNBC is disproportionately prevalent in African-ancestry women, comparing with women of White descent [9–14]. Moreover, TNBC in African-ancestry women also presents higher mortality rates [12, 15, 16], a younger age of onset [6, 14, 17–19], and a faster and more aggressive tumor development and relapse after chemotherapy [19, 20], independently of other risk factors for BC [21]. Although factors for such disparity may include advanced disease stage at diagnosis, socioeconomic status, and lack of access to treatment [22], biological disparities have been suggested as one of the main factors involved in the ancestry-associated BC discrepancy [22–24]. Although studies have investigated ancestry-associated differences in TNBC patients concerning gene and protein expression [25, 26], somatic alterations [13, 26–28], and metabolic processes [29, 30], implications in TNBC development and clinical setting are not yet clarified.

To identify ancestry-associated molecular differences between African-ancestry and White TNBC patients, taking into consideration patients menopause status and stage of the disease, with potential impact in patients’ prognosis and treatment, we analyzed differential gene expression profile using The Cancer Genome Atlas (TCGA) data, the most used database in BC ancestry-associated studies [11, 26, 28, 31–34]. To ascertain if the differences identified in patients from TCGA are transposed to clinical samples we further selected four differentially expressed candidates, EGFR, Myc, Bcl2 and β-Catenin, which were then validated through immunohistochemistry in samples gathered in a Portuguese hospital. Intriguingly, despite the geographic and cultural differences, the immunohistochemistry results match the bioinformatic analysis.

## Methods

### Patient populations

From the publicly available TCGA’s BC project data, “TCGA-BRCA”, we identified 52 African-ancestry and 89 White patients diagnosed with primary TNBC, who had undergone RNA-sequencing. Patients were diagnosed between 1998 and 2013, and clinical information was submitted to TCGA between August 2010 and March 2015. RNA-sequencing and clinical data were retrieved from TCGA using *TCGAbiolinks* package [35, 36] in R environment (version 3.6.0), as of March 6, 2019. To independently validate the results obtained with the TCGA cohort, formalin-fixed paraffin-embedded (FFPE) TNBC samples were collected from Hospital Professor Doutor Fernando Fonseca (HFF)’s patients. 12 African-ancestry and 11 White TNBC patients diagnosed between January 2012 and August 2018 comprised the validation cohort.

### Differential Gene Expression Analysis

Differential gene expression analysis (DGEA) was performed with *edgeR* [37], version 3.26.8, using generalized linear model (gml)-*edgeR* pipeline [38] and a false discovery rate (FDR) cutoff of ≤0.05. We applied a filtering threshold of at least 10 counts-per-million in two or more patients/libraries in each DGEA. The list of differential expressed genes (DEG) was further narrowed down using *glmTreat()* function by testing whether the differential expression was significantly above a log_2_-fold-change (FC) of 1.2. This strategy enabled us to focus on more biological meaningful genes. Comparisons between different lists of differentially expressed genes was performed with *UpSetR* package [39]. Gene set enrichment analysis (GSEA) and Ingenuity Pathway Analysis (IPA, QIAGEN Inc., https://www.qiagenbioinformatics.com/products/ingenuity-pathway-analysis) software were used to identify key regulators and pathways.

For GSEA, *GAGE* “Generally Applicable Gene-set Enrichment” package [40] was used to bring Kyoto Encyclopedia of Genes and Genomes (KEGG) gene sets into the R environment. *edgeR*’s *fry()* function, based on the ROAST method [41], was used to perform GSEA. GSEA included gene sets from the following collections: signal transduction, cellular community - eukaryotes and cell motility, cell growth and death, cancer: overview, where the BC gene set was included, as well as the drug resistance - antineoplastic collection, women-specific endocrine system collection gene sets, and gene sets related to immuno oncology, from the immune system collection. Gene sets were considered enriched when FDR ≤0.05, taking into consideration both up- and downregulated genes in the set.

IPAs’ core analysis, using the complete lists of genes and respective FC and FDR obtained from DGEA, allowed the identification of relevant altered networks, as well as the central network regulators, which are DEG with more direct and indirect connections in its network, being potential drivers for downstream ancestry-associated biological differences observed in TNBC patients. Ingenuity Knowledge Base (Genes only) was used as reference. Only networks with a score above 30 were considered. A threshold of FDR≤0.05 was applied to highlight DEG in a network.

### Immunohistochemistry

Immunohistochemistry (IHC) was performed to validate candidate DEG potentially involved in the ancestry-associated discrepancy of TNBC, using FFPE samples. After deparaffinization, endogenous peroxidase blocking and antigen retrieval, samples were incubated 1h with primary antibody (Primary antibodies and respective dilutions are listed in the Supplementary Table S1), followed by 1h incubation with the secondary antibody (Peroxidase IgG Fraction Monoclonal Mouse Anti-Rabbit IgG, light chain specific, code 211-032-171, Jackson ImmunoResearch). DAB (Bright-DAB Substract Kit, IL ImmunoLogic, VWR) was used for protein detection. Images were acquired using Axio Imager Z2 microscope (Zeiss). Protein expression was evaluated using Fiji ImageJ [42] software and a semiquantitative approach to assign an H-score [43, 44]. The intensity of protein expression was measured using the scale 0-3, 0 being negative and 3 being very high expression. The H-Score was calculated by multiplying the intensity value with its area. Wilcoxon-Mann-Whitney test was used to compare mean H-score of each protein between African-ancestry and White patients using the R function *wilcox.test()* and GraphPad Prism 6.01 software.

### Statistical analysis

Wilcoxon-Mann-Whitney test was used to compare age of diagnosis and survival between groups of patients. Pearson’s Chi-squared test with Yates’ continuity correction was used to test for statistically significant differences between the percentage of patients of each ancestry regarding staging and menopause status, using *chisq.test()* R function. *survival* [45] and *survminer* [46] packages were used to draw and test for differences regarding the time of survival between African-ancestry and White TNBC patients according to staging and menopause status. P-values<0.05 were considered statistically significant.

## Results

### TNBC incidence and patients’ survival

Distribution of clinicopathologic features from 940 African-American (AA) and White BC patients from “TCGA-BRCA” project are depicted in Table 1. AAs are more likely to develop TNBC than their White counterparts (28.42% vs 11.89%, p<0.0001), and are diagnosed with BC at earlier age (56.29 vs 58.61 years, p=0.037). Differences in patients’ age, pathological stage and menopause status at TNBC diagnosis were not statistically significant, as well as months of follow-up, vital status and time to death. Supplementary - Patients file describes patients’ characteristics.

**Table 1.** Description of TCGA’s African-American and White breast cancer patients.^a^

|  | Black or African-American<br>(n=183) | White<br>(n=757) | P-value |
| --- | --- | --- | --- |
| Age at diagnosis, mean (IQR) | 56.29 (47 - 67) | 58.61 (49 - 67) | 0.037<sup>b</sup> |
| Tumor type, n (%) |  |  |  |
| **TNBC** | 52 (28.42) | 90 (11.89) | <0.0001 |
| HER2 | 7 (3.83) | 22 (2.91) | 0.684 |
| Hormone receptor | 112 (61.20) | 589 (77.81) | <0.0001 |
| Indeterminate or not evaluated | 12 (6.56) | 56 (7.40) |  |
| Age at diagnosis of **TNBC**, mean (IQR) | 55.87 (48 - 62.75) | 54.23 (46.25 - 62) | 0.780<sup>b</sup> |
| Stage at **TNBC** diagnosis, n (%) |  |  |  |
| I | 9 (17.31) | 19 (21.11) | 0.741 |
| II | 32 (61.54) | 56 (62.22) | >0.999 |
| III | 10 (19.23) | 11 (12.22) | 0.375 |
| IV | 1 (1.92) | 1 (1.11) | NA |
| Indeterminate or not evaluated | - | 3 (3.33) | NA |
| Menopause status at **TNBC** diagnosis, n (%) |  |  |  |
| Pre (<6 months since LMP, no prior bilateral ovariectomy and not on estrogen replacement) | 9 (17.31) | 29 (32.22) | 0.082 |
| Peri (6-12 months since last menstrual period) | 3 (5.77) | 1 (1.11) | NA |
| Post (prior bilateral ovariectomy or >12 months since LMP with no prior hysterectomy) | 32 (61.54) | 55 (61.11) | 0.999 |
| Indeterminate or not evaluated | 8 (15.48) | 5 (5.56) | NA |
| Follow-up **TNBC** patients, months (IQR)<sup>c</sup>, | 36.83 (14.70 - 50.77) | 28.79 (5.78 - 44.92) | 0.932<sup>b</sup> |
| Vital Status of **TNBC** patients, n (%) |  |  | 0.883 |
| Alive | 45 (86.54) | 80 (88.89) |  |
| Dead | 7 (13.46) | 10 (11.11) | NA |
| Time to death of **TNBC** patients, months (IQR) | 30.75 (25.13 - 36.80) | 42.73 (12.43 - 58.94) | 0.743<sup>b</sup> |

<sup>a</sup> Some values do not sum to heading totals because of missing data.<sup>b</sup> Wilcoxon-Mann-Whitney test. Otherwise, the comparisons are by Pearson's Chi-squared test.<sup>c</sup> Months of follow-up after initial diagnosis.
NA – Not available
IQR – Interquartile range
LMP – Last menstrual period

TNBC patients were then divided according to their ancestry and controlled for the menopause status and pathological stage of TNBC at diagnosis, since these variables may contribute to molecular disparities in TNBC biology. The survival probability for each group was analyzed up to 12 years follow-up after initial diagnosis (Fig. 1). Although not statistically significant, AA patients tended to have a faster and higher mortality than White patients when all patients with TNBC or postmenopausal patients were considered (Fig 1a and c), in line with what have been reported. Interestingly, this tendency is statistically significant within premenopausal and stage II TNBC patients (Fig. 1b and e, log-rank p=0.019 and 0.0038, respectively).

**Fig. 1.**
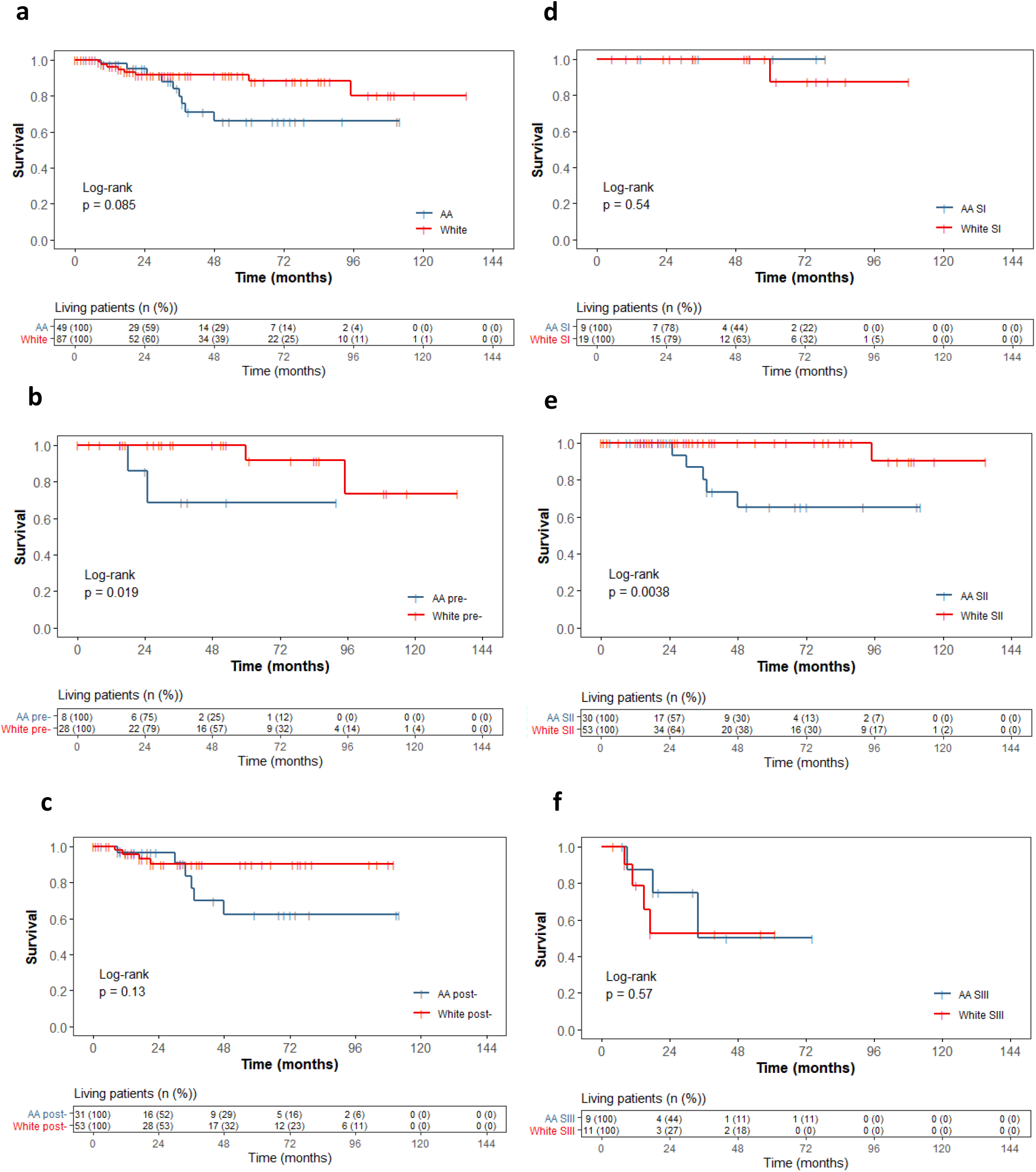
Kaplan-Meier survival curves of TNBC cases stratified by ancestry up to 12 years follow-up after initial diagnosis. **a** All TNBC patients. **b** Premenopausal (pre-) TNBC patients. **c** Postmenopausal (post-) TNBC patients. **d** Stage I (SI) TNBC patients. **e** Stage II (SII) TNBC patients. **f** Stage III (SIII) TNBC patients. Blue survival curves correspond to African-American patients (AA), red curves show White patients. Values do not sum to heading totals because of missing follow-up information.

### Differential gene expression analysis

Differential expression between groups of patients was firstly explored through multi-dimensional scaling (MDS) plots (Fig. S1). In accordance with previous studies [26, 31], and despite some outlier patients, gene expression profiles dissimilarities are higher between TNBC patients and patients with other subtypes of BC (Hormone receptor [HR+] and HER2), than between patients diagnosed with HR+ and HER2 BCs (Fig. S1a). Also, non-TNBC patients (HR+ and HER2) have a more dissimilar expression profile from TNBC patients, regardless of patients’ ancestry (Fig. S1b). The MDS plot does not show a clear differential expression profile between AA and White TNBC patients (Fig. S1c). However, ancestry-associated discrepancies at transcriptomic level were previously reported [11, 25, 26, 31, 47, 48], motivating further analysis using more refined methodologies.

In order to identify gene expression profile differences between AA and White TNBC patients, which may explain the less favorable survival probability of AAs, the following groups were subjected to DGEA: 1) All AA vs all White patients; 2) AA vs White patients with matched menopause status; 3) AA vs White patients with matched TNBC stage; 4) AA vs White patients with matched menopause status and TNBC stage.

Figure 2 shows the number of significantly DEG (FDR≤0.05) in AA patients comparing with matching White patients obtained in the different DGEA, as well as the number of matching DEG between different groups of patients. DGEA controlling for stage II patients resulted in the higher number of DEG (1776) although it is a subset of the full dataset. This result highlights the importance of the disease staging in explaining the differences between the two patients groups. DGEA including all TNBC patients recovers 1122 DEG and DGEA controlling for post-menopause, with a similar number of patients than the stage II comparison, shows 718 DEGs. Furthermore, DGEA controlling for stage III TNBC had threefold the DEG than DGEA controlling for stage I (94 vs 30, respectively), even though the first has less patients included. Although studies consistently report an earlier age of TNBC onset in African-ancestry patients [6, 14, 18, 19], our DGEA results suggest that, except for stage I, gene expression profile distance is greater when controlling for disease’s pathological stage than when controlling for menopause status. Complete DGEA results are listed in Supplementary - DGEA file. Matching DEG obtained in different DGEA can be consulted in Supplementary - Matching DEG file.

**Fig. 2.**
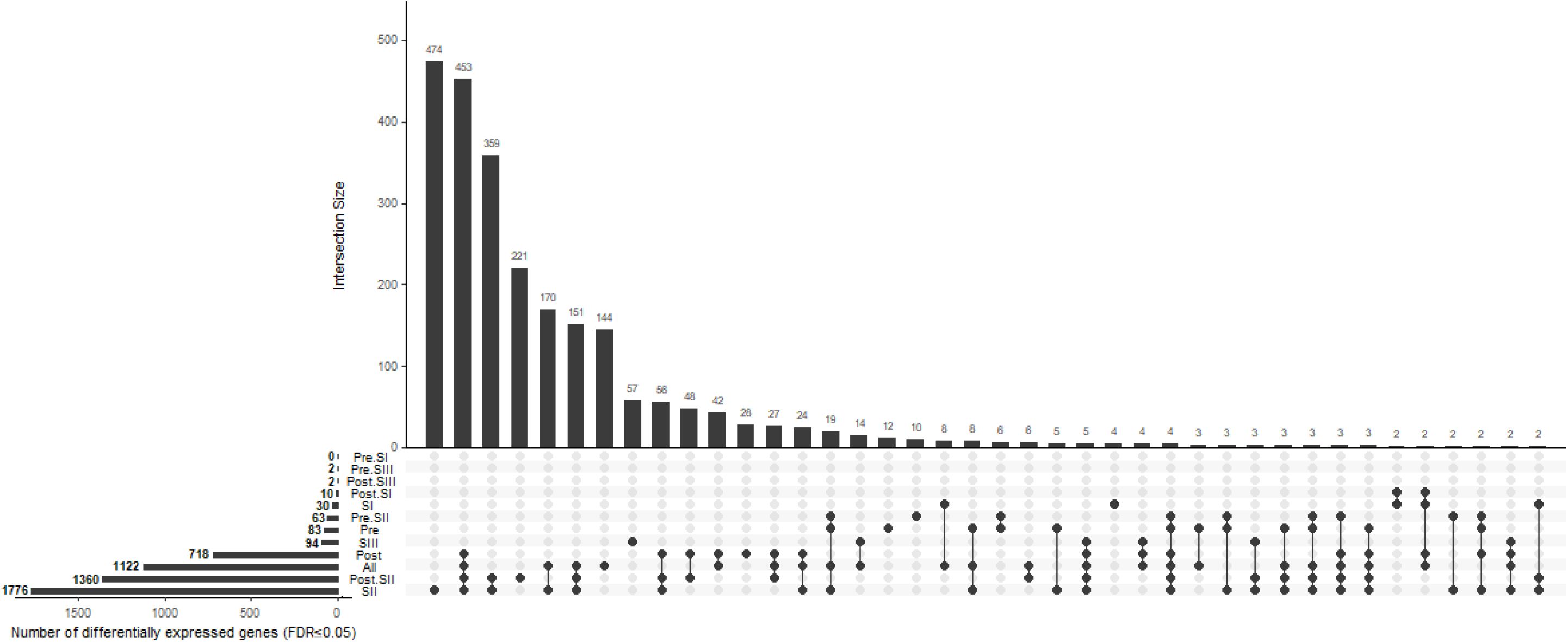
UpSet plot of the differentially expressed genes (FDR≤0.05) idenfied in TNBC AA patients comparing with TNBC White patients, obtained in the different differential gene expression analyses (DGEA). The intersection size represents the number of matching genes. *All* DGEA considering all TNBC patients (52 AA vs 89 White). *Pre* DGEA considering premenopausal patients (9 AA vs 29 White). *Post* DGEA considering postmenopausal patients (32 AA vs 57 White). *SI* DGEA considering stage I disease patients (9 AA vs 19 White). *SII* DGEA considering stage II disease patients (32 AA vs 59 White). *SIII* DGEA considering stage III disease patients (10 AA vs 12 White). *PreSI* DGEA considering premenopausal stage I disease patients (2 AA vs 7 White). *PreSII* DGEA considering premenopausal stage II disease patients (5 AA vs 17 White). *PreSIII* DGEA considering premenopausal stage III disease patients (2 AA vs 4 White). *PostSI* DGEA considering postmenopausal stage I disease patients (3 AA vs 11 White). *PostSII* DGEA considering postmenopausal stage II disease patients (22 AA vs 36 White). *PostSIII* DGEA considering postmenopausal stage III disease patients (7 AA vs 6 White). *AA* African-American.

Additionally, DGEA with RNA-sequencing data from normal-adjacent BC tissue from AA and White patients (76 DEG, 6 AA vs 105 White) confirmed that the large majority of DEG identified in DGEA are likely associated with how TNBC disease progresses in the two patients populations, differences that are not visible in normal (non-TNBC) tissue. In fact, only two genes - DEG *CXCL10* and *LTB* - were found as to be simultaneously differentially expressed between AA and White patients in both TNBC and normal tissue.

DGEA results showed that there are indeed gene expression ancestry-associated differences to justify TNBC’s epidemiological and clinical observations. Thus, we proceeded to refine DGEA results, in order to find potential leading molecules involved in the ancestry-associated TNBC discrepancy.

### Gene set enrichment analysis and Ingenuity Pathway Analysis

GSEAs were performed to contextualize ancestry-associated biological differences of AA and White TNBC patients taking in account their menopausal status and disease’s pathological stage. Here, only the groups that showed significantly enriched gene sets (FDR.Mixed≤0.05, Fig. 3) are presented, namely all TNBC, postmenopausal (independently of the disease stage), stage II (independently of menopausal status) and postmenopausal and stage IIs. Complete GSEA results can be consulted in Supplementary - GSEA file.

**Fig. 3.**
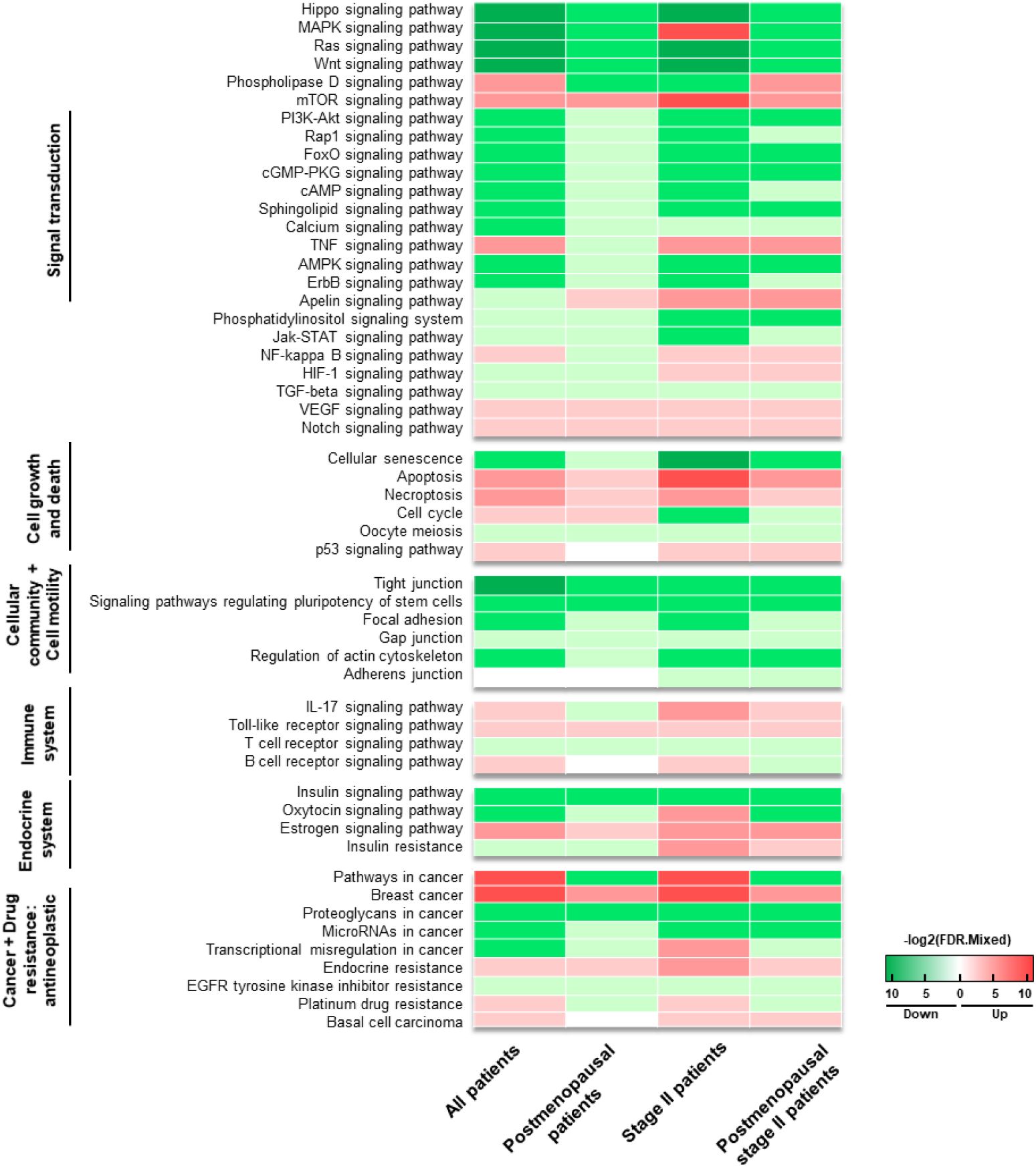
Enriched gene sets in AA patients. Each column corresponds to a group of patients, in the following order: all TNBC patients, postmenopausal patients, stage II disease patients and postmenopausal stage II disease patients. Green color - negative net direction (Down). Red color - positive net direction (Up). White color – not statistically significant (FDR.Mixed≥0.05). Color intensity is proportional to −log_2_(FDR.Mixed) value.

Considering the uniformity of gene sets’ FC net direction among all four groups of patients, as well as their statistical significance, we highlight Hippo signaling pathway as the most consistently downregulated enriched gene set, and the cellular community and cell motility as the most downregulated gene set collection in all groups of patients. Interestingly, we also highlight the BC gene set as being consistently upregulated in all the presented groups, and MAPK signaling pathway gene set as being upregulated specifically in the group with stage II disease patients.

According to the literature, downregulation of Hippo pathway components (Fig. S2a), including *LATS1*, may promote the transcription of genes involved in cell proliferation and competition, cell death inhibition, epithelial-to-mesenchymal transition, tumor metastasis, and tumorigenesis [49–54]. Cellular community and cell motility gene sets (Fig. S2b-g) are consistently downregulated, suggesting that the integrity of cell-cell contacts and actin cytoskeleton organization, fundamental in epithelial tissue homeostasis, are impaired [55, 56], promoting tumor cells dissociation and subsequent metastasis [57]. We also highlight that, BC gene set (Fig. S2h), which includes, among others, *EGFR*, *MYC* or *CTNNB1* genes, is upregulated in AA patients comparing with White patients, suggesting that these women have a higher activation of components and processes involved in BC development, which may be translated into TNBC’s faster progression and aggressive behavior. Interestingly, MAPK signaling pathway gene set (Fig. S2i) is exclusively upregulated in stage II disease AA patients. MAPK cascades regulate a wide variety of cellular processes, including proliferation, differentiation, apoptosis and stress responses, playing a crucial role in the survival and development of cancer cells [58]. Thus, DEG involved in MAPK pathway exclusively identified in stage II AA patients, including *NTRK1*, *GADD45C*, *MAPK12*, *TRADD* or *FGFR2*, may contribute to the observed worst prognosis of AA TNBC patients when comparing with matching White patients.

To identify potential driving molecules involved in the ancestry-associated TNBC discrepancy, we also performed IPA’s core analysis with focus on network design, using the complete lists of genes obtained through DGEA for each group, except for pre- and postmenopausal stage I and III disease, due to their low number of DEG (Figure 4). Networks in IPA allow the visualization of the interactions between molecules. Thus, highly interconnected networks are likely to represent significant biological functions. Complete lists of networks and respective central regulators, scores, molecules involved, and top diseases and functions, according to patient’s menopause status and/or staging, are listed in Supplementary - Networks file.

**Fig. 4.**
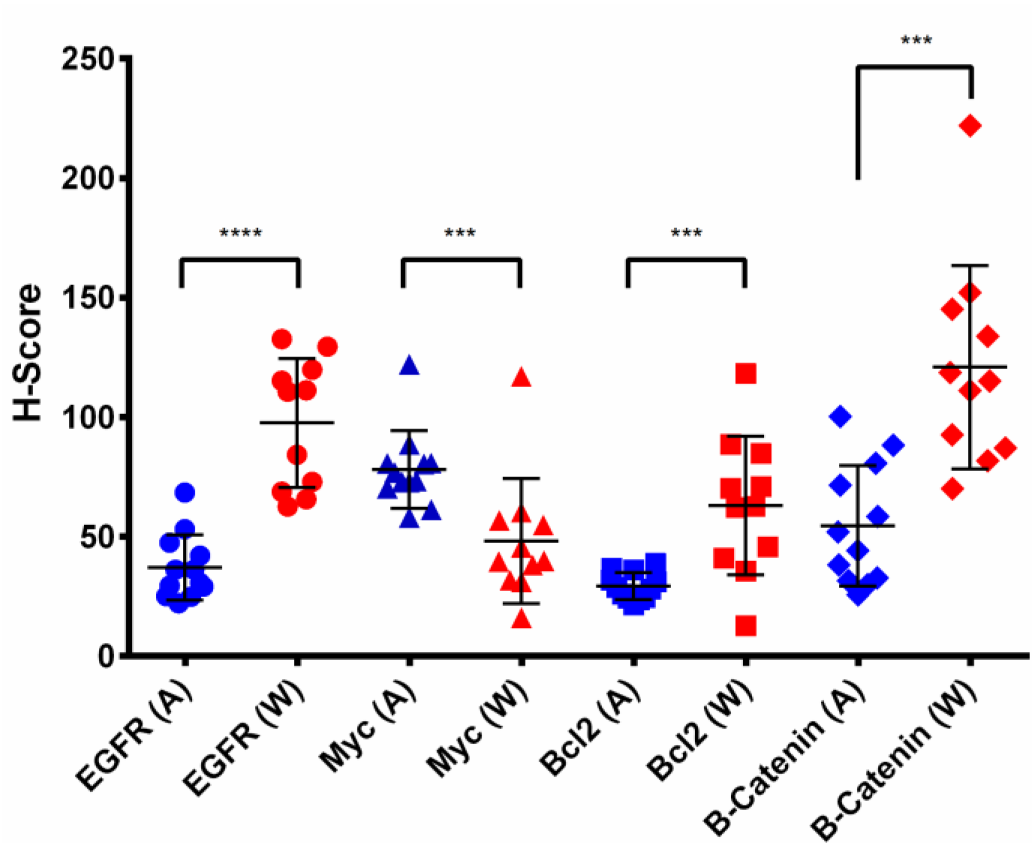
Differences in H-score between African-ancestry (n=12) and White (n=11) patients are significantly different for EGFR (p=5.92E-06), Myc (p=4.01E-04), Bcl2 (p=9.81E-04) and β-Catenin (p=1.44E-04) expression and have the same net direction as the results obtained in DGEA. A/Blue - African-ancestry patients. W/Red - White patients. Circles - EGFR expression. Triangles - Myc expression. Squares - Bcl2 staining. Diamonds - β-catenin expression.

Here we highlighted EGFR as central network regulator (Fig. S3) due to its high score and consistency in the groups with all TNBC patients and patients presenting post-menopause, stage II disease, or both. Specifically, *EGFR* is downregulated in AA patients, comparing with White patients, in these four groups (log_2_FC=−2.238, FDR=1.08E-04; log_2_FC=−2.911, FDR=9.41E-05; log_2_FC=−2.775, FDR=9.85E-05; and log_2_FC=−3.739, FDR=4.46E-05, respectively). Furthermore, EGFR is involved in enriched gene sets from the Signal transduction collection, including MAPK, Ras, Phospholipase D, PI3K-Akt, Rap1, FoxO, calcium, ErbB, Jak-STAT and HIF-1 signaling pathways; in gene sets from the Cancer and Drug resistance: antineoplastic collections, including BC, pathways, proteoglycans and microRNAs in cancer and EGFR tyrosine kinase inhibitor resistance; in Cellular community and Cell motility collections, including regulation of actin cytoskeleton, focal adhesion and gap junction gene sets; and in oxytocin and estrogen signaling pathways gene sets from the Endocrine system collection.

We also highlighted Myc as a central network regulator (Fig. S4) due to its score and consistency in the groups with all TNBC patients, postmenopausal and postmenopausal stage II disease patients. Indeed, *MYC* is upregulated in AA patients, regarding White patients, from these groups (log_2_FC=0.754, FDR=2.70E-02; log_2_FC=1.051, FDR=1.13E-02; log_2_FC=1.279, and FDR=5.16E-03, respectively). Also, Myc is involved in enriched gene sets from the Signal transduction collection, including Hippo, MAPK, Wnt, PI3K-Akt, ErbB, Jak-STAT and TGF-beta signaling pathways; Cancer collection’s BC, pathways, proteoglycan, microRNAs and transcriptional misregulation in cancer gene sets; in Cell growth and death collection’s cellular senescence and cell cycle gene sets; and signaling pathways regulating pluripotency of stem cells gene set from Cellular community collection.

Finally, we highlight Bcl2 as a highly scored central network regulator (Fig. S5) specifically downregulated in premenopausal AA patients, relatively to White patients (log_2_FC=−2,043, FDR=3,49E-02). No enriched gene sets were observed with FDR.mixed≤0.05, possible due to the small set of significant DEG observed in this group (83 DEG, Table 3). Interestingly, when comparing with normal-adjacent BC tissue from AA patients, *BCL2* is downregulated in TNBC patients, considering all menopause status (log_2_FC=−2.19, FDR=1.10E-04, Supplementary - DGEA), suggesting that *BCL2* is downregulated throughout TNBC development in younger AA patients.

Thus, EGFR, Myc and Bcl2 were selected to be further investigated through IHC in a collection of TNBC samples obtained from a Portuguese hospital. This procedure was performed to ascertain if the differential protein and mRNA expression would match, even though these populations may present geographic and cultural differences from the original populations sampled in TCGA. Additionally, we investigated β-catenin, encoded by *CTNNB1*, due to the well-described role of Wnt/β-catenin signaling in BC and TNBC tumorigenesis [59–61] and worse prognosis [59, 62–64]. Also, *CTNNB1* is downregulated in postmenopausal stage II disease AA patients (log_2_FC=−0.935, FDR=1.63E-02), and is involved in the following enriched gene sets: Hippo and Wnt signaling pathways, from the Signal transduction collection; BC gene set; and Cellular community collection’s signaling pathways regulating pluripotency of stem cells gene set.

### Validation of EGFR, Myc, Bcl2 and β-catenin protein expression

African-ancestry patients from the validation cohort (Table 2) were diagnosed with TNBC at a significantly younger age, comparing with White patients (47.75 vs 64.82 years, p=0.007). Accordingly, premenopausal TNBC cases in the African-ancestry patients are in a significantly higher proportion, comparing with White patients (58.33% vs 0, p=0.010). None of the other clinical variables had statistically significant differences between the two populations of patients in the validation cohort.

**Table 2.** Description of the validation cohort.

|  | African-ancestry<br>(n=12) | White<br>(n=11) | P-value |
| --- | --- | --- | --- |
| Age at diagnosis of **TNBC**, mean (IQR) | 47.75 (37 - 54.25) | 64.82 (58 - 70) | 0.007<sup>a</sup> |
| Stage at **TNBC** diagnosis, n (%) |  |  |  |
| I | 1 (8.33) | 4 (36.36) | 0.262 |
| II | 4 (33.33) | 2 (18.18) | 0.725 |
| III | 2 (16.67) | 2 (18.18) | >0.999 |
| IV | 4 (33.33) | 2 (18.18) | 0.725 |
| Indeterminate or not evaluated | 2 (16.67) | 1 (9.1) | NA |
| Menopause status at **TNBC** diagnosis, n (%) |  |  |  |
| Pre (<6 months since LMP, no prior bilateral ovariectomy and not on estrogen replacement) | 7 (58.33) | 0 | 0.010 |
| Post (prior bilateral ovariectomy or >12 months since LMP with no prior hysterectomy) | 3 (25) | 10 (90.9) | 0.006 |
| Indeterminate or not evaluated | 2 (16.67) | 1 (9.1) | NA |
| Follow-up **TNBC** patients, mean (IQR)<sup>b</sup> | 43 (21 - 58) | 57 (45 - 73) | 0.131 |
| Vital Status of **TNBC** patients, n (%) |  |  | >0.999 |
| Alive | 10 (83.33) | 10 (90.9) | NA |
| Dead | 2 (16.67) | 1 (9.1) | NA |
| Time to death of **TNBC** patients, months | 22 - 33 | 53 | 0.667 |

<sup>a</sup> Wilcoxon-Mann-Whitney test. Otherwise, the comparisons are by Fisher exact test.
<sup>b</sup> Months of follow-up after initial diagnosis.
NA – Not available
IQR – Interquartile range
LMP – Last menstrual period

Validation of gene expression through isolation of RNA from FFPE samples followed by quantitative RT-PCR was not possible due to the poor of quality of paraffin-embedded tissue samples, which did not allowed good RNA extraction. Thus, IHC was performed to validate the selected candidates, showing that EGFR, Myc, Bcl2 and β-Catenin proteins are differentially expressed between African-ancestry and White patients (p≤0.05, Fig. 4) and the net direction of protein expression intensity matches with DGEA results. Therefore, these observations corroborate that while Myc is upregulated in African-ancestry patients, EGFR, Bcl2 and CTNNB1/β-Catenin are downregulated, in comparison with White patients.

Examples of positive and low/negative staining between African-ancestry and White patients for each of the tested proteins are shown in Fig. 5. Of note the fact that all African-ancestry patients had low/negative Bcl2 expression (Fig. 5c).

**Fig. 5.**
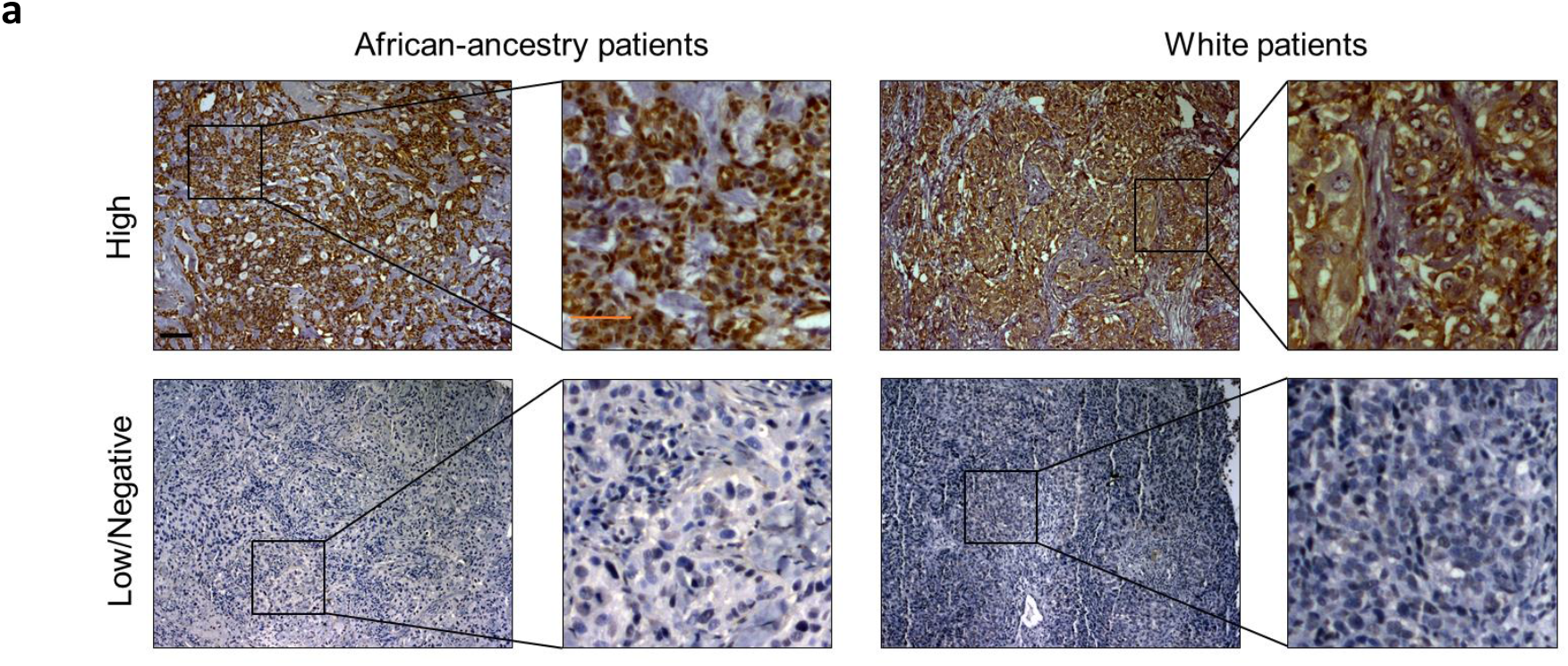
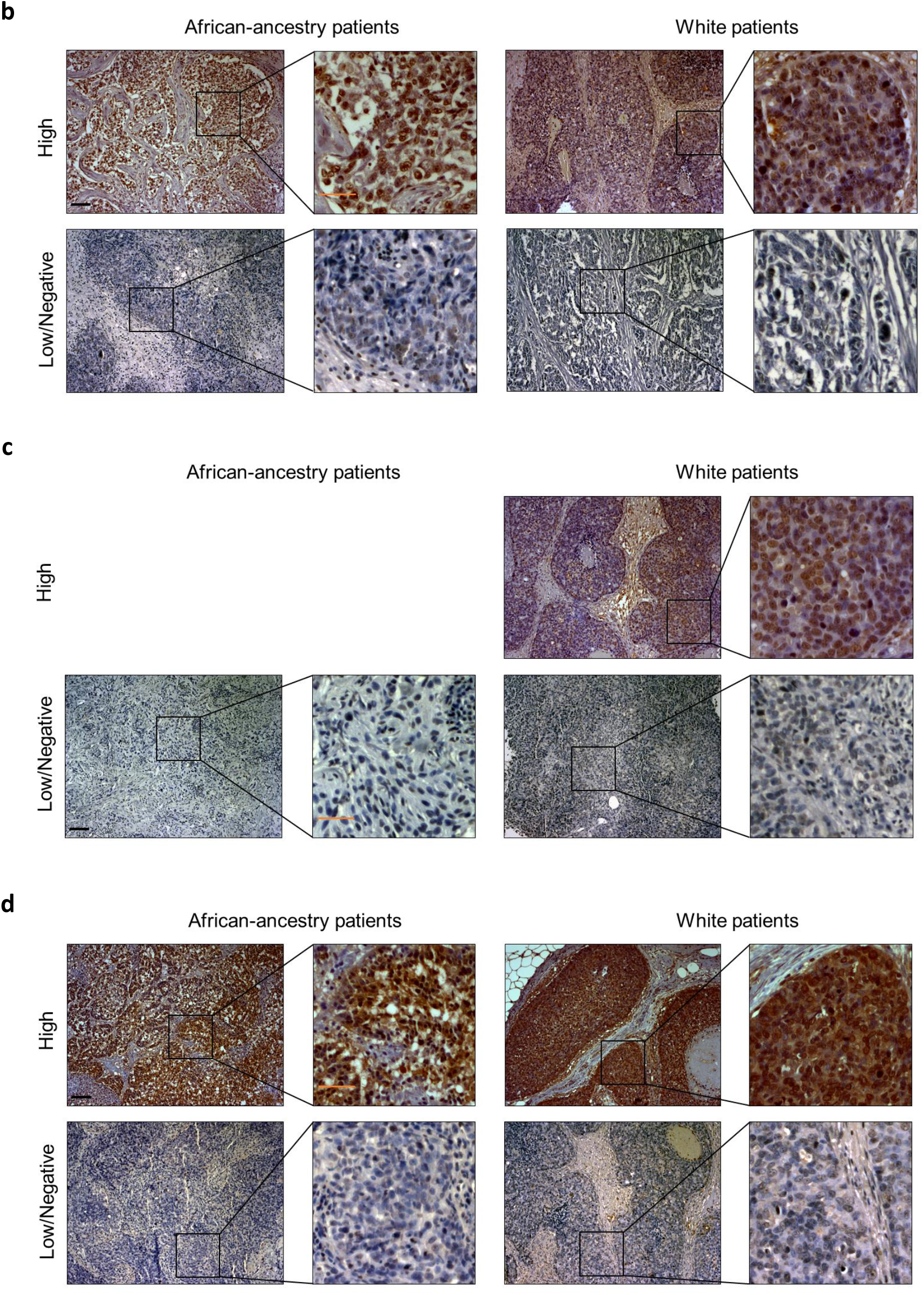
Immunohistochemistry staining of formalin-fixed paraffin embed TNBC tissue from African-ancestry patients (left) and White patients (right) with **a.** anti-EGFR IgG, **b.** anti-Myc IgG, **c.** anti-Bcl2 IgG and **d.** anti-β-Catenin IgG. Brown color indicates positive reactivity (top), showing protein expression. Black scale bar - 100 μm. Orange scale bar - 50 μm.

## Discussion

Clinical differences between TNBC incidence and behavior in African-ancestry and White women are expected to heavily influence differences in the transcriptomic profile and regulation of pathways associated with tumorigenesis. Identification of ancestry-associated markers should contribute to understand more clearly the clinical features of this disease in both populations. Additionally, these studies may also contribute for patients’ prognosis and establishment of more effective targeted therapies, potentially closing the widening mortality gap between these populations. Our study is one of the few to systematically examine ancestry-associated differences at transcriptomic level in TNBC patients, with the particularity of considering patients’ menopause status and/or pathological stage of the disease. Furthermore, the subsequent validation of candidate markers through IHC, in sociocultural and geographically distinct patients, make our findings more robust.

“TCGA-BRCA” epidemiological study confirmed a higher TNBC incidence in AA patients, corroborating previous TCGA-based studies [11, 26, 31, 33, 34], and a decreased survival probability in AAs diagnosed specifically at pre-menopause and at stage II disease. Also, although TNBC has a discrepant age of onset between African-ancestry and White patients [6, 14, 18], which was also observed in the validation cohort (Table 2), DGEA results suggest that differences in ancestry-associated transcriptomic programs in TNBC patients are more pronounced when associated with disease staging than with patients’ age or hormonal differences caused by menopause, as observed by the higher number of DEG identified when controlling for stage II disease, even though it has a lower number of patients than other DGEAs. Accordingly, stage III disease patients DGEA resulted in a number of DEG threefold as higher than DGEA between stage I patients, despite having a similar number of patients.

To identify altered pathways and biological processes in TNBC AA patients, comparing with their White counterparts, GSEA was performed. Among the enriched gene sets observed in the groups with all TNBC patients or controlling for post-menopause condition, stage II disease, or both, we highlighted those with a consistent net direction among groups, including Hippo signaling pathway, the most consistently downregulated enriched gene set, potentially promoting tumorigenesis; gene sets from the cellular community and cell motility gene collections, also downregulated, suggesting loss of cell-cell contact integrity and potential metastasis formation; BC gene set, consistently upregulated in all of those AA groups, even though White patients also have TNBC; and MAPK signaling pathway, being upregulated specifically in stage II disease AA patients, suggesting that this pathway may contribute to the observed survival discrepancies.

IPA’s network design, through central regulator identification and network scoring, also contributed to the selection of the following candidate ancestry-associated markers, which were further investigated through IHC: EGFR, Myc and Bcl2. Furthermore, we also analyzed β-Catenin expression due to its known involvement in BC and TNBC tumorigenesis [59–61].

EGFR is a transmembrane receptor for members of the epidermal growth factor family, reportedly promoting cell proliferation, motility, and survival via activation of various downstream signaling pathways, including MAPK and PI3K-AKT-mTOR [65]. EGFR is frequently reported as overexpressed in several cancers, including TNBC, being associated with poor prognosis, shorter disease-free survival and TNBC aggressiveness [66–71]. Curiously, in our study, EGFR gene and protein expression were downregulated in African-ancestry patients. Few studies reported EGFR under-expression as being associated with BC progression or metastasis formation [72, 73], and patients’ poor prognosis [74], however, Kreike’s study [74] suggested that EGFR expression in BC patients has a nonlinear relation with disease outcome, with both lower and higher expression associated with poor prognosis and worse outcome compared to intermediate expression levels. The precise oncogenic role of EGFR downregulation remains puzzlingly elusive, particularly in TNBC development and prognosis in African-ancestry patients.

Myc transcription factor, regulating up to 15% of human genes, is involved in cell growth, proliferation, metabolism, differentiation, stress pathways, drug resistance and apoptosis, sustaining growth of many types of cancers [75–77]. Oncogenic Myc expression is increased in TNBC, as compared to other BC subtypes [78–81], contributing to chemoresistance and poor prognosis [80, 81]. Myc, has been stated as being overexpressed in TNBC in studies mostly composed by White patients or cell lines [78–81]. Here, we observed that African-ancestry patients have an even higher expression of Myc. In line with this observation, a TCGA-based study reported that TNBC AA patients have a higher proportion of *MYC*/8q24.21 amplification, comparing with White patients (30.9% vs 20.4%), independently of other variables [26].

Apoptosis regulator Bcl2 reportedly acts by promoting cell survival instead of driving cell proliferation. Impaired apoptosis preserves preneoplastic cells and those bearing mutations that ablate their need for trophic factors, potentially facilitating metastasis to distant sites [82]. Previously, two studies suggested Bcl2 expression as a favorable prognostic marker for HR-positive BCs [83, 84], and other study as an independent indicator of favorable prognosis for all types of early-stage BC, including TNBC [85]. Nonetheless, the effects of *BCL2*/Bcl2 low expression in cancer, and its implications in patients’ prognosis and treatment, are poorly explored. We observed downregulation of *BCL2* expression specifically in premenopausal AA patients, comparing with White patients, as well as a significantly low/negative protein expression in African-ancestry patients, mostly premenopausal. Interestingly, the fact that *BCL2* is significantly downregulated in TNBC comparing with normal-adjacent BC tissue in AAs (Supplementary - DGEA), suggests that this downregulation may contribute to TNBC development in these premenopausal African-ancestry patients. Our observation is in line with the Abdel-Fatah study [86], which has shown that a negative expression of Bcl2 in early primary-TNBCs was significantly associated with high proliferation and an increased risk of death and recurrence. Furthermore, they also observed that although Bcl2-negative patients improved survival and disease-free survival upon adjuvant-anthracycline combination-chemotherapy treatment, when these patients do not achieve a pathological complete response, their prognosis is worse [86]. Thus, overall, *BCL2*/Bcl2 expression may be a promising prognosis marker for African-ancestry patients with TNBC, particularly at younger ages.

Finally, β-Catenin plays a central role in canonical Wnt signaling pathway and as an intermediate in other signal transduction pathways, including PI3K-AKT pathway. The role and impact of *CTNNB1*/β-catenin expression and misregulation in BC have been massively studied [64, 87–95], following the crescent number of studies about Wnt/β-catenin signaling targeting as a therapeutic target in many tumors, especially colorectal carcinomas [96–99]. In TNBC, enrichment of canonical Wnt signal pathway and β-catenin expression is reportedly required for TNBC development by controlling numerous tumor-associated properties, including migration, stemness, anchorage-independent growth and chemosensitivity [61], being associated with poor clinical outcome [59, 60, 62–64]. Interestedly, in this study, we observed that African-ancestry TNBC patients express less *CTNNB1*/β-catenin comparing with their White counterparts, despite the usual worst prognosis in these population. Therefore, the current paradigm regarding *CTNNB1*/β-catenin role in TNBC should not be generalized.

Remarkably, even though DGEA and IHC were completely different approaches, performed based on USA patients’ RNA-sequencing data and in samples from Portuguese hospitals, respectively, gene and protein differential expression matched. AA population is predominantly of Niger-Kordofanian/western Africa ancestry [100, 101], while 8/12 African-ancestry patients followed in Portugal are from western Africa countries, namely Cape Verde, São Tomé and Guinea-Bissau (Supplementary - Patients file). These observations suggest that at least patients with western Africa ancestry, besides having an higher incidence of TNBC [13, 14], also share some TNBC-associated features, distinct from White patients.

Our results reinforce that TNBC is a highly heterogeneous disease with ancestry-associated biological differences that should also be considered in addition to the molecular differences between normal vs BC or TNBC vs other BC. Although African-ancestry patients are less participative in clinical trials [102–104], our study show how important it is to include a diverse group of participants in tumor marker, cell-based and clinical trial studies. Importantly, ancestry assumptions should not be considered biological guideposts, as commonly defined racial/populational groups are genetically heterogeneous and lack clear-cut genetic boundaries [100, 105]. Thus, although we suggest EGFR, Myc, Bcl2 and β-catenin expression as ancestry-associated markers in African-ancestry patients with TNBC, we cannot exclude the possibility of African-ancestry patients having those chromosomic regions, or the chromosomic regions of genes regulating these markers, of other ancestries [101]. Furthermore, our findings should be complemented by a validation in a larger group of patients, as well as further functional studies resourcing to cell lines and animal models.

In conclusion, although self-described ancestry should not substitute the specific disease’s etiology for each individual patient, predictions based on patients’ self-described ancestry in clinical settings may be helpful in predicting patients’ prognosis and proper treatments, being an easy and inexpensive guiding method.

## Abbreviations

AA: African-American
BC: Breast cancer
DEG: Differential expressed genes
DGEA: Differential gene expression analysis
FC: Fold-change
FDR: False discovery rate
FFPE: Formalin-fixed paraffin-embedded
GSEA: Gene set enrichment analysis
HER2: Human Epidermal growth factor Receptor 2
HFF: Hospital Professor Doutor Fernando Fonseca
HR: Hormone receptor
IHC: Immunohistochemistry
IPA: Ingenuity Pathway Analysis
KEGG: Kyoto Encyclopedia of Genes and Genomes
MDS: Multi-dimensional scaling
TCGA: The Cancer Genome Atlas
TNBC: Triple-negative breast cancer

## Acknowledgements

We are grateful to all HFF’s BC patients that agreed to participate in this study, as well as the Pathology department for the assistance regarding sample collection. We also thank Ramiro Magno and Joana Xavier, at Universidade do Algarve, for the teachings involving R Programming Environment. We acknowledge the help with IPA from Bruno Alexandre at Instituto de Biologia Experimental e Tecnológica. We also appreciate Ana Farinho and Telmo Pereira for the assistance regarding the IHC methodology and microscopy, respectively.

## Funding

This work was supported by national Portuguese funding through Fundação para a Ciência e Tecnologia - PD/BD/114053/2015, iNOVA4Health – UIDB/04462/2020 and ALGARVE2020 POCI-01-0145-FEDER-022184-“GenomePT” grant.

## Compliance with ethical standards Conflict of interest

Authors declare no conflicts of interest.

## Ethical approval

The Review Boards at NMS|FCM and HFF approved the use of patient samples in this study. All participants gave informed consent and this study was performed in accordance with the ethical standards outlined in the 1964 Declaration of Helsinki. Informed consent was obtained from all individual participants included in TCGA. Information about TCGA can be found at https://www.cancer.gov/aboutnci/organization/ccg/research/structural-genomics/tcga

## Notes

### Competing Interest Statement

The authors have declared no competing interest.

## References

1. Collett K, Stefansson IM, Eide J, et al (2005) A basal epithelial phenotype is more frequent in interval breast cancers compared with screen detected tumors. Cancer Epidemiol Biomarkers Prev 14:1108–1112. https://doi.org/10.1016/S1043-321X(05)80267-X

2. Seewaldt VL, Scott V (2007) Rapid progression of basal-type breast cancer. N Engl J Med 356:2015. https://doi.org/10.1056/NEJMicm063760

3. Garrido-Castro AC, Lin NU, Polyak K (2019) Insights into molecular classifications of triple-negative breast cancer: Improving patient selection for treatment. Cancer Discov 9:176–198. https://doi.org/10.1158/2159-8290.CD-18-1177

4. Saraiva DP, Guadalupe Cabral M, Jacinto A, Braga S (2017) How many diseases is triple negative breast cancer: The protagonism of the immune microenvironment. ESMO Open 2:. https://doi.org/10.1136/esmoopen-2017-000208

5. Vona-davis L, Rose DP, Hazard H, et al (2008) Triple-Negative Breast Cancer and Obesity in a Rural Appalachian Population. 17:3319–3325. https://doi.org/10.1158/1055-9965.EPI-08-0544

6. Carey LA, Perou CM, Livasy CA, et al (2006) Race, Breast Cancer Subtypes, and Survival in the Carolina Breast Cancer Study. JAMA 295:2492–2502

7. Rakha EA, El-Rehim DA, Paish C, et al (2006) Basal phenotype identifies a poor prognostic subgroup of breast cancer of clinical importance. Eur J Cancer 42:3149–3156. https://doi.org/10.1016/j.ejca.2006.08.015

8. Rakha EA, El-Sayed ME, Green AR, et al (2007) Prognostic markers in triple-negative breast cancer. Cancer 109:25–32. https://doi.org/10.1002/cncr.22381

9. Stark A, Kleer CG, Martin I, et al (2010) African ancestry and higher prevalence of triple-negative breast cancer. Cancer. Cancer 116:4926–4932. https://doi.org/10.1002/cncr.25276.African

10. Boyle P (2012) Triple-negative breast cancer: Epidemiological considerations and recommendations. Ann Oncol 23:vi7–vi12. https://doi.org/10.1093/annonc/mds187

11. Keenan T, Moy B, Mroz EA, et al (2015) Comparison of the genomic landscape between primary breast cancer in African American versus white women and the association of racial differences with tumor recurrence. J Clin Oncol 33:3621–3627. https://doi.org/10.1200/JCO.2015.62.2126

12. DeSantis CE, Ma J, Gaudet MM, et al (2019) Breast cancer statistics, 2019. CA Cancer J Clin 69:438–451. https://doi.org/10.3322/caac.21583

13. Newman LA, Jenkins B, Chen Y, et al (2019) Hereditary Susceptibility for Triple Negative Breast Cancer Associated with Western Sub-Saharan African Ancestry: Results from an International Surgical Breast Cancer Collaborative. Ann Surg 270:484–492. https://doi.org/10.1097/SLA.0000000000003459

14. Huo D, Ikpatt F, Khramtsov A, et al (2009) Population differences in breast cancer: Survey in indigenous african women reveals over-representation of triple-negative breast cancer. J Clin Oncol 27:4515–4521. https://doi.org/10.1200/JCO.2008.19.6873

15. Honório M, Guerra-Pereira N, Silva J, et al (2016) Decreased Survival in African Patients with Triple Negative Breast Cancer. J Palliat Care Med 6:4. https://doi.org/10.4172/2165-7386.1000270

16. Bauer KR, Brown M, Cress RD, et al (2007) Descriptive analysis of estrogen receptor (ER)-negative, progesterone receptor (PR)-negative, and HER2-negative invasive breast cancer, the so-called triple-negative phenotype: A population-based study from the California Cancer Registry. Cancer 109:1721–1728. https://doi.org/10.1002/cncr.22618

17. Lund MJ, Trivers KF, Porter PL, et al (2009) Race and triple negative threats to breast cancer survival: A population-based study in Atlanta, GA. Breast Cancer Res Treat 113:357–370. https://doi.org/10.1007/s10549-008-9926-3

18. Zaky S, Lund M, May K, et al (2009) The Triple Threat of Recurrence after Breast Conserving Therapy: Race, Receptor Status and Age. Cancer Res 69:6045. https://doi.org/10.1158/0008-5472.sabcs-09-6045

19. Copson E, Maishman T, Gerty S, et al (2014) Ethnicity and outcome of young breast cancer patients in the United Kingdom: The POSH study. Br J Cancer 110:230–241. https://doi.org/10.1038/bjc.2013.650

20. Frasci G, Comella P, Rinaldo M, et al (2009) Preoperative weekly cisplatin-epirubicin-paclitaxel with G-CSF support in triple-negative large operable breast cancer. Ann Oncol 20:1185–1192. https://doi.org/10.1093/annonc/mdn748

21. Stead LA, Lash TL, Sobieraj JE, et al (2009) Triple-negative breast cancers are increased in black women regardless of age or body mass index. Breast Cancer Res 11:1–10. https://doi.org/10.1186/bcr2242

22. Newman LA (2017) Breast Cancer Disparities: Socioeconomic Factors versus Biology. Ann Surg Oncol 24:2869–2875. https://doi.org/10.1245/s10434-017-5977-1

23. Grunda JM, Steg AD, He Q, et al (2012) Differential expression of breast cancer-associated genes between stage- and age-matched tumor specimens from African- and Caucasian-American Women diagnosed with breast cancer. BMC Res Notes 5:. https://doi.org/10.1186/1756-0500-5-248

24. Siddharth S, Sharma D (2018) Racial disparity and triple-negative breast cancer in African-American women: A multifaceted affair between obesity, biology, and socioeconomic determinants. Cancers (Basel) 10:1–19. https://doi.org/10.3390/cancers10120514

25. Lindner R, Sullivan C, Offor O, et al (2013) Molecular phenotypes in triple negative breast cancer from African American patients suggest targets for therapy. PLoS One 8:. https://doi.org/10.1371/journal.pone.0071915

26. Huo D, Hu H, Rhie SK, et al (2017) Comparison of Breast Cancer Molecular Features and Survival by African and European Ancestry in The Cancer Genome Atlas. JAMA Oncol 3:1654–1662. https://doi.org/10.1001/jamaoncol.2017.0595

27. Shimelis H, LaDuca H, Hu C, et al (2018) Triple-negative breast cancer risk genes identified by multigene hereditary cancer panel testing. J Natl Cancer Inst 110:855–862. https://doi.org/10.1093/jnci/djy106

28. Omilian AR, Wei L, Chen C, et al (2020) Somatic mutations of triple ‑ negative breast cancer : a comparison between Black and White women. Breast Cancer Res Treat. https://doi.org/10.1007/s10549-020-05693-4

29. Pelicano H, Zhang W, Liu J, et al (2014) Mitochondrial dysfunction in some triple-negative breast cancer cell lines: Role of mTOR pathway and therapeutic potential. Breast Cancer Res 16:1–16. https://doi.org/10.1186/s13058-014-0434-6

30. Tayyari F, Gowda GAN, Olopade OF, et al (2018) Metabolic profiles of triple-negative and luminal A breast cancer subtypes in African-American identify key metabolic differences. Oncotarget 9:11677–11690. https://doi.org/10.18632/oncotarget.24433

31. Stewart PA, Luks J, Roycik MD, et al (2013) Differentially expressed transcripts and dysregulated signaling pathways and networks in African American breast cancer. PLoS One 8:1–13. https://doi.org/10.1371/journal.pone.0082460

32. Polak P, Kim J, Braunstein LZ, et al (2017) A mutational signature reveals alterations underlying deficient homologous recombination repair in breast cancer. Nat Genet 49:1476–1486. https://doi.org/10.1038/ng.3934

33. Ademuyiwa FO, Tao Y, Luo J, et al (2017) Differences in the mutational landscape of triple-negative breast cancer in African Americans and Caucasians. Breast Cancer Res Treat 161:491–499. https://doi.org/10.1007/s10549-016-4062-y

34. O’Meara T, Safonov A, Casadevall D, et al (2019) Immune microenvironment of triple-negative breast cancer in African-American and Caucasian women. Breast Cancer Res Treat 175:247–259. https://doi.org/10.1007/s10549-019-05156-5

35. Colaprico A, Silva TC, Olsen C, et al (2016) TCGAbiolinks: An R/Bioconductor package for integrative analysis of TCGA data. Nucleic Acids Res 44:e71. https://doi.org/10.1093/nar/gkv1507

36. Mounir M, Lucchetta M, Silva TC, et al (2019) New functionalities in the TCGAbiolinks package for the study and integration of cancer data from GDC and GTEX. PLoS Comput Biol 15:. https://doi.org/10.1371/journal.pcbi.1006701

37. Robinson MD, McCarthy DJ, Smyth GK (2009) edgeR: A Bioconductor package for differential expression analysis of digital gene expression data. Bioinformatics 26:139–140. https://doi.org/10.1093/bioinformatics/btp616

38. McCarthy DJ, Chen Y, Smyth GK (2012) Differential expression analysis of multifactor RNA-Seq experiments with respect to biological variation. Nucleic Acids Res 40:4288–4297. https://doi.org/10.1093/nar/gks042

39. Conway JR, Lex A, Gehlenborg N (2017) Genome analysis UpSetR : an R package for the visualization of intersecting sets and their properties. 33:2938–2940. https://doi.org/10.1093/bioinformatics/btx364

40. Luo W, Friedman MS, Shedden K, et al (2009) GAGE: Generally applicable gene set enrichment for pathway analysis. BMC Bioinformatics 10:1–17. https://doi.org/10.1186/1471-2105-10-161

41. Wu D, Lim E, Vaillant F, et al (2010) ROAST: Rotation gene set tests for complex microarray experiments. Bioinformatics 26:2176–2182. https://doi.org/10.1093/bioinformatics/btq401

42. Schindelin J, Arganda-Carreras I, Frise E, et al (2012) Fiji: An open-source platform for biological-image analysis. Nat Methods 9:676–682. https://doi.org/10.1038/nmeth.2019

43. Ishibashi H, Suzuki T, Suzuki S, et al (2003) Sex steroid hormone receptors in human thymoma. J Clin Endocrinol Metab 88:2309–2317. https://doi.org/10.1210/jc.2002-021353

44. John T, Liu G, Tsao MS (2009) Overview of molecular testing in non-small-cell lung cancer: Mutational analysis, gene copy number, protein expression and other biomarkers of EGFR for the prediction of response to tyrosine kinase inhibitors. Oncogene 28:S14–S23. https://doi.org/10.1038/onc.2009.197

45. Therneau T (1999) A package for survival analysis in S. Citeseer 1–83

46. Kassambara A, Kosinski M, Biecek P, Fabian S (2019) survminer: Drawing Survival Curves using “ggplot2”

47. Sugita B, Gill M, Mahajan A, et al (2016) Differentially expressed miRNAs in triple negative breast cancer between African-American and non-Hispanic white women. Oncotarget 7:79274–79291. https://doi.org/10.18632/oncotarget.13024

48. Chang CS, Kitamura E, Johnson J, et al (2018) Genomic analysis of racial differences in triple negative breast cancer. Genomics 1–14. https://doi.org/10.1016/j.ygeno.2018.10.010

49. Piccolo S, Dupont S, Cordenonsi M (2014) The biology of YAP/TAZ: Hippo signaling and beyond. Physiol Rev 94:1287–1312. https://doi.org/10.1152/physrev.00005.2014

50. Zanconato F, Cordenonsi M, Piccolo S (2016) YAP/TAZ at the Roots of Cancer. Cancer Cell 29:783–803. https://doi.org/10.1016/j.ccell.2016.05.005

51. Zhao B, Li L, Lei Q, Guan K-L (2010) The Hippo–YAP pathway in organ size control and tumorigenesis: an updated version. Gen Devel 24:862–874. https://doi.org/10.1101/gad.1909210.The

52. Lei Q-Y, Zhang H, Zhao B, et al (2008) TAZ Promotes Cell Proliferation and Epithelial-Mesenchymal Transition and Is Inhibited by the Hippo Pathway. Mol Cell Biol 28:2426–2436. https://doi.org/10.1128/mcb.01874-07

53. Harvey KF, Zhang X, Thomas DM (2013) The Hippo pathway and human cancer. Nat Rev Cancer 13:246–257. https://doi.org/10.1038/nrc3458

54. Marti P, Stein C, Blumer T, et al (2015) YAP promotes proliferation, chemoresistance, and angiogenesis in human cholangiocarcinoma through TEAD transcription factors. Hepatology 62:1497–1510. https://doi.org/10.1002/hep.27992

55. Cavallaro U, Christofori G (2004) Multitasking in tumor progression: Signaling functions of cell adhesion molecules. Ann N Y Acad Sci 1014:58–66. https://doi.org/10.1196/annals.1294.006

56. Bhat AA, Uppada S, Achkar IW, et al (2019) Tight junction proteins and signaling pathways in cancer and inflammation: A functional crosstalk. Front Physiol 10:1–19. https://doi.org/10.3389/fphys.2018.01942

57. Bogenrieder T, Herlyn M (2003) Axis of evil: Molecular mechanisms of cancer metastasis. Oncogene 22:6524–6536. https://doi.org/10.1038/sj.onc.1206757

58. Guo YANJUN, Pan WEIWEI, Liu SB, et al (2020) ERK / MAPK signalling pathway and tumorigenesis (Review). 1997–2007. https://doi.org/10.3892/etm.2020.8454

59. Dey N, Young B, Abramovitz M, et al (2013) Differential Activation of Wnt-β-Catenin Pathway in Triple Negative Breast Cancer Increases MMP7 in a PTEN Dependent Manner. PLoS One 8:1–17. https://doi.org/10.1371/journal.pone.0077425

60. Dey N, Barwick BG, Moreno CS, et al (2013) Wnt signaling in triple negative breast cancer is associated with metastasis. BMC Cancer 13:. https://doi.org/10.1186/1471-2407-13-537

61. Xu J, Prosperi JR, Choudhury N, et al (2015) Β-Catenin Is Required for the Tumorigenic Behavior of Triple-Negative Breast Cancer Cells. PLoS One 10:1–11. https://doi.org/10.1371/journal.pone.0117097

62. Khramtsov AI, Khramtsova GF, Tretiakova M, et al (2010) Wnt/β-catenin pathway activation is enriched in basal-like breast cancers and predicts poor outcome. Am J Pathol 176:2911–2920. https://doi.org/10.2353/ajpath.2010.091125

63. Geyer FC, Lacroix-Triki M, Savage K, et al (2011) Β-Catenin pathway activation in breast cancer is associated with triple-negative phenotype but not with CTNNB1 mutation. Mod Pathol 24:209–231. https://doi.org/10.1038/modpathol.2010.205

64. López-Knowles E, Zardawi SJ, McNeil CM, et al (2010) Cytoplasmic localization of â-catenin is a marker of poor outcome in breast cancer patients. Cancer Epidemiol Biomarkers Prev 19:301–309. https://doi.org/10.1158/1055-9965.EPI-09-0741

65. Yarden Y (2001) The EGFR family and its ligands in human cancer: Signalling mechanisms and therapeutic opportunities. Eur J Cancer 37:3–8. https://doi.org/10.1016/s0959-8049(01)00230-1

66. Nielsen TO, Hsu FD, Jensen K, et al (2004) Immunohistochemical and clinical characterization of the basal-like subtype of invasive breast carcinoma. Clin Cancer Res 10:5367–5374. https://doi.org/10.1158/1078-0432.CCR-04-0220

67. Viale G, Rotmensz N, Maisonneuve P, et al (2009) Invasive ductal carcinoma of the breast with the “triple-negative” phenotype: Prognostic implications of EGFR immunoreactivity. Breast Cancer Res Treat 116:317–328. https://doi.org/10.1007/s10549-008-0206-z

68. Park HS, Jang MH, Kim EJ, et al (2014) High EGFR gene copy number predicts poor outcome in triple-negative breast cancer. Mod Pathol 27:1212–1222. https://doi.org/10.1038/modpathol.2013.251

69. Nakai K, Hung M-C, Yamaguchi H (2016) A perspective on anti-EGFR therapies targeting triple-negative breast cancer. Am J Cancer Res 6:1609–23

70. Ali R, Wendt MK (2017) The paradoxical functions of EGFR during breast cancer progression. Signal Transduct Target Ther 2:1–7. https://doi.org/10.1038/sigtrans.2016.42

71. Foidart P, Yip C, Radermacher J, et al (2019) Expression of MT4-MMP, EGFR, and Rb in triple-negative breast cancer strongly sensitizes tumors to erlotinib and palbociclib combination therapy. Clin Cancer Res 25:1838–1850. https://doi.org/10.1158/1078-0432.CCR-18-1880

72. Choong LY, Lim S, Loh MCS, et al (2007) Progressive loss of epidermal growth factor receptor in a subpopulation of breast cancers: Implications in target-directed therapeutics. Mol Cancer Ther 6:2828–2842. https://doi.org/10.1158/1535-7163.MCT-06-0809

73. Wendt MK, Williams WK, Pascuzzi PE, et al (2015) The Antitumorigenic Function of EGFR in Metastatic Breast Cancer is Regulated by Expression of Mig6. Neoplasia (United States) 17:124–133. https://doi.org/10.1016/j.neo.2014.11.009

74. Kreike B, Hart G, Bartelink H, Van De Vijver MJ (2010) Analysis of breast cancer related gene expression using natural splines and the Cox proportional hazard model to identify prognostic associations. Breast Cancer Res Treat 122:711–720. https://doi.org/10.1007/s10549-009-0588-6

75. Camarda R, Zhou AY, Kohnz RA, et al (2016) Inhibition of fatty acid oxidation as a therapy for MYC-overexpressing triple-negative breast cancer. Nat Med 22:427–432. https://doi.org/10.1038/nm.4055

76. Fallah Y, Brundage J, Allegakoen P, Shajahan-Haq AN (2017) MYC-Driven pathways in breast cancer subtypes. Biomolecules 7:1–6. https://doi.org/10.3390/biom7030053

77. Dang C V., O’Donnell KA, Zeller KI, et al (2006) The c-Myc target gene network. Semin Cancer Biol 16:253–264. https://doi.org/10.1016/j.semcancer.2006.07.014

78. Horiuchi D, Kusdra L, Huskey NE, et al (2012) MYC pathway activation in triple-negative breast cancer is synthetic lethal with CDK inhibition. J Exp Med 209:679–696. https://doi.org/10.1084/jem.20111512

79. Koboldt DC, Fulton RS, McLellan MD, et al (2012) Comprehensive molecular portraits of human breast tumours. Nature 490:61–70. https://doi.org/10.1038/nature11412

80. Lee K min, Giltnane JM, Balko JM, et al (2017) MYC and MCL1 Cooperatively Promote Chemotherapy-Resistant Breast Cancer Stem Cells via Regulation of Mitochondrial Oxidative Phosphorylation. Cell Metab 26:633–647.e7. https://doi.org/10.1016/j.cmet.2017.09.009

81. Carey JPW, Karakas C, Bui T, et al (2018) Synthetic lethality of PARP inhibitors in combination with MYC blockade is independent of BRCA status in triple-negative breast cancer. Cancer Res 78:742–757. https://doi.org/10.1158/0008-5472.CAN-17-1494

82. Adams JM, Cory S (2007) The Bcl-2 apoptotic switch in cancer development and therapy. Oncogene 26:1324–1337. https://doi.org/10.1038/sj.onc.1210220

83. Eom YH, Kim HS, Lee A, et al (2016) BCL2 as a subtype-specific prognostic marker for breast cancer. J Breast Cancer 19:252–260. https://doi.org/10.4048/jbc.2016.19.3.252

84. Choi JE, Kang SH, Lee SJ, Bae YK (2014) Prognostic significance of Bcl-2 expression in non-basal triple-negative breast cancer patients treated with anthracycline-based chemotherapy. Tumor Biol 35:12255–12263. https://doi.org/10.1007/s13277-014-2534-4

85. Dawson S, Makretsov N, Blows FM, et al (2010) BCL2 in breast cancer : a favourable prognostic marker across molecular subtypes and independent of adjuvant therapy received. Br J Cancer 668–675. https://doi.org/10.1038/sj.bjc.6605736

86. Abdel-Fatah TMA, Perry C, Dickinson P, et al (2013) Bcl2 is an independent prognostic marker of triple negative breast cancer (TNBC) and predicts response to anthracycline combination (ATC) chemotherapy (CT) in adjuvant and neoadjuvant settings. Ann Oncol 24:2801–2807. https://doi.org/10.1093/annonc/mdt277

87. Lin SY, Xia W, Wang JC, et al (2000) β-catenin, a novel prognostic marker for breast cancer: Its roles in cyclin D1 expression and cancer progression. Proc Natl Acad Sci U S A 97:4262–4266. https://doi.org/10.1073/pnas.060025397

88. Prasad CP, Gupta SD, Rath G, Ralhan R (2008) Wnt signaling pathway in invasive ductal carcinoma of the breast: Relationship between β-catenin, disheveled and cyclin D1 expression. Oncology 73:112–117. https://doi.org/10.1159/000120999

89. Dolled-Filhart M, McCabe A, Giltnane J, et al (2006) Quantitative in situ analysis of β-catenin expression in breast cancer shows decreased expression is associated with poor outcome. Cancer Res 66:5487–5494. https://doi.org/10.1158/0008-5472.CAN-06-0100

90. Fanelli MA, Montt-Guevara M, Diblasi AM, et al (2008) P-Cadherin and β-catenin are useful prognostic markers in breast cancer patients; β-catenin interacts with heat shock protein Hsp27. Cell Stress Chaperones 13:207–220. https://doi.org/10.1007/s12192-007-0007-z

91. Bànkfalvi À, Terpe HJ, Breukelmann D, et al (1999) Immunophenotypic and prognostic analysis of E-cadherin and β-catenin expression during breast carcinogenesis and tumour progression: A comparative study with CD44. Histopathology 34:25–34. https://doi.org/10.1046/j.1365-2559.1999.00540.x

92. Chung GG, Zerkowski MP, Ocal IT, et al (2004) β-Catenin and p53 Analyses of a Breast Carcinoma Tissue Microarray. Cancer 100:2084–2092. https://doi.org/10.1002/cncr.20232

93. Dillon DA, D’Aquila T, Reynolds AB, et al (1998) The expression of p120ctn protein in breast cancer is independent of α- and β-catenin and E-cadherin. Am J Pathol 152:75–82

94. Nakopoulou, Gakiopoulou, Keramopoulos, et al (2000) C-Met Tyrosine Kinase Receptor Expression Is Associated With Abnormal Β-Catenin Expression and Favourable Prognostic Factors in Invasive Breast Carcinoma. Histopathology 36:313–325. https://doi.org/10.1046/j.1365-2559.2000.00847.x

95. Karayiannakis AJ, Nakopoulou L, Gakiopoulou H, et al (2001) Expression patterns of β-catenin in in situ and invasive breast cancer. Eur J Surg Oncol 27:31–36. https://doi.org/10.1053/ejso.1999.1017

96. Shang S, Hua F, Hu ZW (2017) The regulation of β-catenin activity and function in cancer: Therapeutic opportunities. Oncotarget 8:33972–33989. https://doi.org/10.18632/oncotarget.15687

97. Cheng X, Xu X, Chen D, et al (2019) Therapeutic potential of targeting the Wnt/β-catenin signaling pathway in colorectal cancer. Biomed Pharmacother 110:473–481. https://doi.org/10.1016/j.biopha.2018.11.082

98. O’Toole SA, Beith JM, Millar EKA, et al (2013) Therapeutic targets in triple negative breast cancer. J Clin Pathol 66:530–542. https://doi.org/10.1136/jclinpath-2012-201361

99. Krishnamurthy N, Kurzrock R (2018) Targeting the Wnt/beta-catenin pathway in cancer: Update on effectors and inhibitors. Cancer Treat Rev 62:50–60. https://doi.org/10.1016/j.ctrv.2017.11.002

100. Tishkoff SA, Reed FA, Friedlaender FR, et al (2009) The Genetic Structure and History of Africans and African Americans. Science (80−) 324:1035–1044. https://doi.org/10.1126/science.1172257.The

101. Bryc K, Auton A, Nelson MR, et al (2010) Genome-wide patterns of population structure and admixture in West Africans and African Americans. Proc Natl Acad Sci U S A 107:786–791. https://doi.org/10.1073/pnas.0909559107

102. Shavers VL, Brown ML (2002) Racial and ethnic disparities in the receipt of cancer treatment. J Natl Cancer Inst 94:334–357. https://doi.org/10.1093/jnci/94.5.334

103. Owens OL, Jackson DD, Thomas TL, et al (2013) African american men’s and women’s perceptions of clinical trials research: Focusing on prostate cancer among a high-risk population in the South. J Health Care Poor Underserved 24:1784–1800. https://doi.org/10.1353/hpu.2013.0187

104. Haynes-Maslow L, Godley P, Dimartino L, et al (2014) African American women’s perceptions of cancer clinical trials. Cancer Med 3:1430–1439. https://doi.org/10.1002/cam4.284

105. Serre D, Pääbo S (2004) Evidence for gradients of human genetic diversity within and among continents. Genome Res 14:1679–1685. https://doi.org/10.1101/gr.2529604

